# Tankyrase interacts with the allosteric site of glucokinase and inhibits its glucose-sensing function in the beta cell

**DOI:** 10.1101/2021.08.08.454510

**Authors:** Nai-Wen Chi, Travis Eisemann, Tsung-Yin J. Yeh, Swati Roy, Tyler J. Chi, Sunny H. Lu, John M. Pascal, Olivia Osborn

**Affiliations:** Division of Endocrinology and Diabetes, Dept. of Medicine, Univ. of California, San Diego, La Jolla, CA, USA; Dept. of Biochemistry and Molecular Biology, Thomas Jefferson University, Philadelphia, PA, USA

## Abstract

Insulin secretion in the pancreatic beta cell is rate-limited by glucokinase (GCK), the glucose sensor that catalyzes the first step of glucose metabolism. GCK consists of two lobes connected by a flexible hinge that allows the kinase to exhibit a spectrum of conformations ranging from the active, closed form to several inactive, less-compact forms. Activating GCK mutations can cause hyperinsulinemia and hypoglycemia in infants. A similar phenotype exhibited by tankyrase (TNKS)-deficient mice prompted us to investigate whether TNKS might modulate the glucose-sensing function of GCK. We found that TNKS colocalizes and directly interacts with GCK. Their interaction is mediated by two ankyrin-repeat clusters (ARC-2 and −5) in TNKS and a tankyrase-binding motif (TBM, aa 63-68) in the GCK hinge. This interaction is conformation sensitive, human GCK variants that cause hyperglycemia (V62M) or hypoglycemia (S64Y) enhance or diminish the interaction respectively, even though they have no impact on TNKS interaction in the context of a GCK peptide (V62M) or a peptide library (S64Y). Moreover, the TNKS-GCK interaction is inhibited by high glucose concentrations, which are known to stabilize GCK in the active (closed, glucose-avid) conformation. Conversely, glucose phosphorylation by GCK *in vitro* is inhibited by TNKS. To study this *in vitro* inhibitory effect in the MIN6 beta cells, we showed that glucose-stimulated insulin secretion is suppressed upon stabilization of the TNKS protein and is conversely enhanced upon TNKS knockdown. Based on these findings as well as by contrasting with hexokinase-2, we propose that TNKS is a physiological GCK inhibitor in pancreatic beta cells that acts by trapping the kinase in the open (inactive) conformation.

## INTRODUCTION

Glucose metabolism in mammals is initiated upon its phosphorylation by hexokinases (HKs). HKs 1, 2, and 3 are ~100-kDa proteins consisting of two structurally superimposable halves, each half in turn consisting of two lobes connected by a hinge [1]. The HK catalytic cycle is often described using the induced-fit model. This model posits that in the absence of glucose, each half of HK assumes an open conformation with the two lobes separated by 12° at the hinge. Upon binding glucose, each half of HKs adopts a closed conformation by apposing the two lobes to form the catalytic center. After catalysis, the hinge opens to return the kinase to its open state [2–4]. HK catalysis exhibits a hyperbolic (Michaelis-Menten) dependence on glucose concentrations with a low K_D_ of 0.3 mM [1].

By contrast, GCK (also known as HK4) is half in size (~50 kDa) and shares ~50% homology with each half of the other hexokinases [3]. It exhibits a sigmoidal dependence on glucose concentrations that is not readily explained using the induced-fit model. Instead, a conformational selection (population-shift) model is often invoked [5, 6]. Specifically, the highly flexible hinge of GCK allows it to exhibit an ensemble of conformations with the two lobes progressively separated by up to 99°. These conformations exist in the absence of glucose, and their glucose affinity decreases progressively from the closed (active) state (K_D_ of 0.2 mM glucose), the open state, the wide-open state, to the super-open state (K_D_ of 30 mM)[7–9]. High levels of glucose progressively shift the population toward the closed (glucose-avid) conformation. Due to the sluggishness of the population shift, the sigmoidal curve of GCK activity rises steeply as glucose concentration increases across the physiological range of 5-10 mM. This unusual feature makes GCK an ideal glucose sensor for the pancreatic beta cells to regulate insulin secretion with an S_0.5_ of 8 mM glucose [10, 11]. Once secreted, insulin acts on multiple organs to lower blood glucose levels, thus closing the glucose-insulin feedback loop.

In intermediary metabolism, glucose stimulates insulin secretion through two regulatory pathways [12]. In the *triggering pathway* that is rate-limited by GCK, glucose metabolism induces K_ATP_ channel closure to trigger membrane depolarization, leading to a cytosolic influx of calcium. In the *amplifying pathway*, calcium’s efficacy on promoting insulin vesicle exocytosis is amplified by multiple metabolic factors including lipid signaling, glucose metabolism, and mitochondrial function [13]. The amplifying pathway is not rate-limited by GCK since it is insensitive to pharmacological GCK activators [14] and it exhibits a hyperbolic instead of a sigmoidal dependence on glucose [15, 16]. Because both pathways are activated by metabolizable secretagogues like glucose, an established strategy to selectively assess the amplifying pathway is to use non-metabolizable secretagogues like KCl and the K_ATP_ channel blocker tolbutamide to activate only the triggering pathway [17].

Consistent with the importance of GCK as the physiological glucose sensor, over 200 inactivating mutations in GCK have been reported to cause hyperglycemia and diabetes in humans [18]. Conversely, a handful of activating mutations in GCK can cause hypoglycemia through excessive insulin secretion [11]. Many of these activating mutations map to connecting loop I (variably defined as aa 47-71 [19] or aa 62-72 [20]) in the GCK hinge, thereby confirming the functional importance of this region [11]. Connecting loop I exhibits dramatic structural flexibility as the enzyme reorganizes its two lobes in response to glucose [3, 19]. It is critical to the enzyme’s sigmoidal dependence on glucose [19], and it provides 4 contact residues (aa 62, 63, 65 and 66) for allosteric GCK activators [3, 21], thus making this region an allosteric site. Within this region, the discovery of a tankyrase-binding motif (TBM), ^63^RSDPEG^68^, is reported herein.

Tankyrase (TNKS) and the closely related TNKS2 are molecular scaffolds that each contains 5 homologous clusters of ankyrin repeats (ARC-1 through 5)[22]. Many of these ARCs interact with diverse proteins bearing the TBM RxxPDG [23]. Proteins recruited to the TNKS scaffold often undergo covalent modification (PARsylation) by the poly-ADP-ribose polymerase (PARP) domain at the TNKS C-terminus. PARsylation typically tags the acceptors, including TNKS itself, toward degradation [24].

TNKS has been implicated in diverse physiological processes including insulin action, energy homeostasis, protein secretion, *wnt* signaling, and telomere stability [25, 26]. In mice, TNKS is highly expressed in the pancreatic beta cells [27]. Our genetic model shows that in TNKS-deficient mice, fasting glucose levels are 11% (0.7 mM) below wild-type controls whereas circulating insulin levels are 70% higher [27]. Since this trait phenocopies infants with hypoglycemia and hyperinsulinemia caused by activating GCK mutations, we set out to explore potential roles of TNKS in regulating the glucose-sensing function of GCK in the beta cells.

## RESULTS

### Colocalization of TNKS and glucokinase

TNKS resides in diverse intracellular locales including the Golgi apparatus just outside the nucleus [28, 29], the centrosome [30], and the mitotic spindle poles [31]. We immunolocalized endogenous GCK in the betacell line MIN6 using 3 independent antibodies. During mitosis, all antibodies localized GCK to the spindle poles (**Fig. 1A**), similar to the known mitotic localization of TNKS [31]. In interphase, the staining of endogenous GCK was weak and inconsistent between the 3 antibodies (data not shown), prompting us to transfect MIN6 cells with GFP-GCK. **Fig. 1B** shows that GFP-GCK resided throughout the cytosol with perinuclear enrichment similar to TNKS.

**Figure 1.**
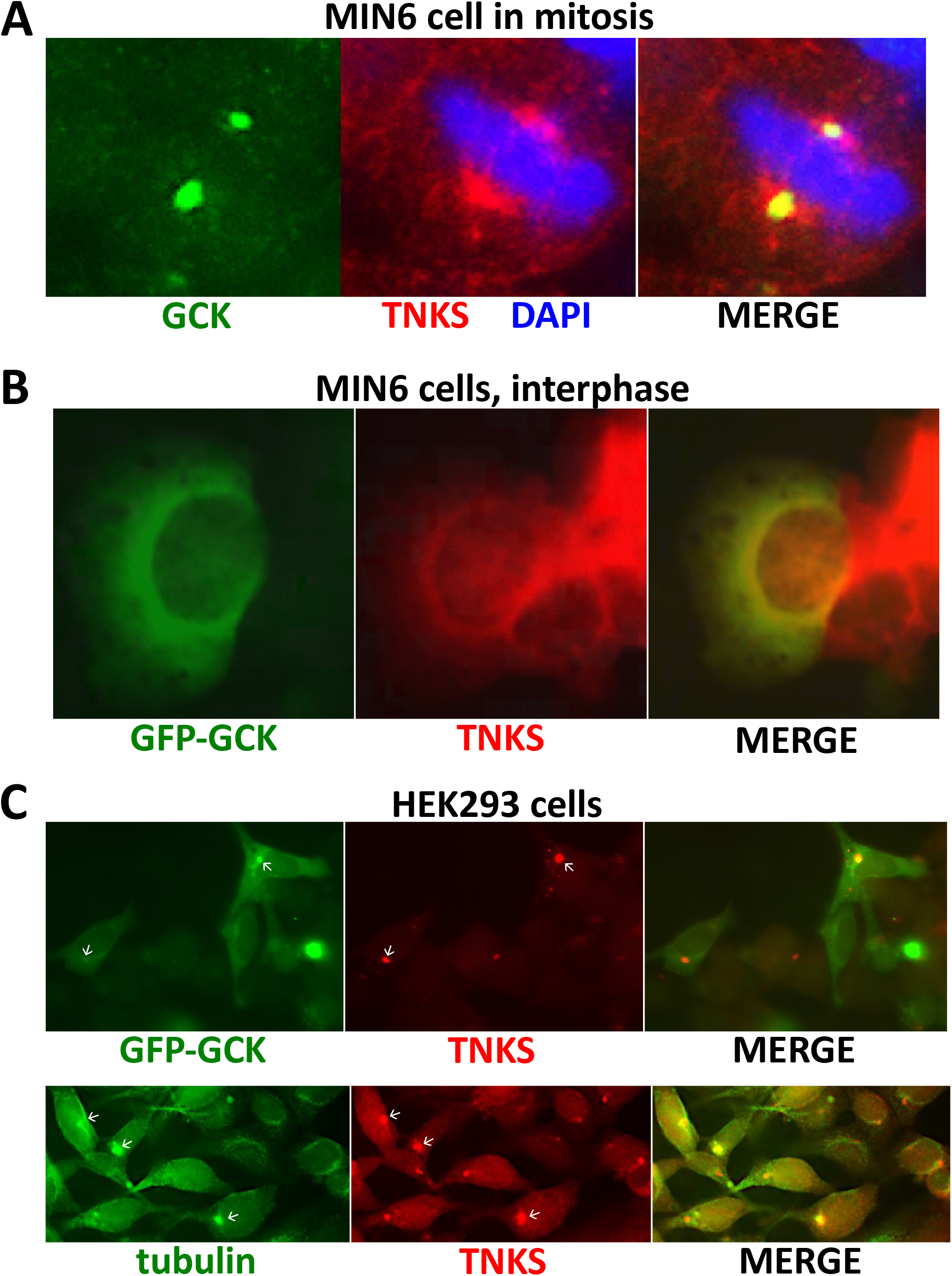
Colocalization of TNKS and glucokinase (GCK) in mitosis and in interphase. **A**. Immunostaining of endogenous GCK (green) and TNKS (red) in mitotic MIN6 cells. A similar spindle polar staining of GCK was observed using two other antibodies as described in Materials and Methods (data not shown). **B**. Localization of transfected GFP-GCK (green) and immunostaining of endogenous TNKS (red) in interphase MIN6 cells. **C.** Localization of transfected GFP-GCK (green) and immunostaining of beta-tubulin (green) and TNKS (red) in HEK239 cells. Arrows indicate centrosomes.

When ectopically expressed in HEK293 cells, the cytosolic distribution of GFP-GCK exhibited distinct enrichment and colocalization with TNKS just outside the nucleus (**Fig. 1C**). The site of their colocalization corresponded to the centrosomes as revealed by tubulin staining (**Fig. 1C**). Since GCK associates with tubulins [32] and colocalizes with TNKS at microtubule-organizing centers (*i.e*., spindle poles and the centrosomes; **Figs. 1A and C**), microtubules seem a plausible platform for GCK and TNKS to colocalize.

### TNKS binds to a TNKS-binding motif (TBM) in the GCK allosteric site

GCK harbors a TNKS-binding motif (RSTPEG; aa 63-68) in its allosteric site, suggesting a potentially conformational-sensitive interaction with TNKS. This was confirmed by using purified GST-GCK fusions to pull down TNKS and TNKS2 from cell lysates (**Fig. 2A**) and using FLAG-tagged TNKS ARC_1-5_ (aa 159-983) to co-precipitate (*myc*)_2_-GCK in transfected cells (**Fig. 2B**). To assess the affinity of the GCK TBM for TNKS, we incubated various concentrations of TNKS ARC_1-5_ with a fluorophore-conjugated GCK peptide (aa 59-74) encompassing the TBM in a fluorescence polarization assay. **Fig. 2C** shows a dissociation constant of ~0.5 μM between the two parties, an affinity comparable to other TBM-containing peptides [33, 34]. To specify the ARCs in TNKS that interacted with GCK, individual ARCs were purified as (His)_6_-SUMO fusions and incubated with GST fusion of the GCK TBM hexapeptide. **Fig. 2D** shows that the GCK TBM pulled down only ARC-2 and −5. Collectively, these findings established TNKS as a novel partner of GCK that uses ARC-2 and −5 to interact with the TBM in the GCK allosteric site.

**Figure 2.**
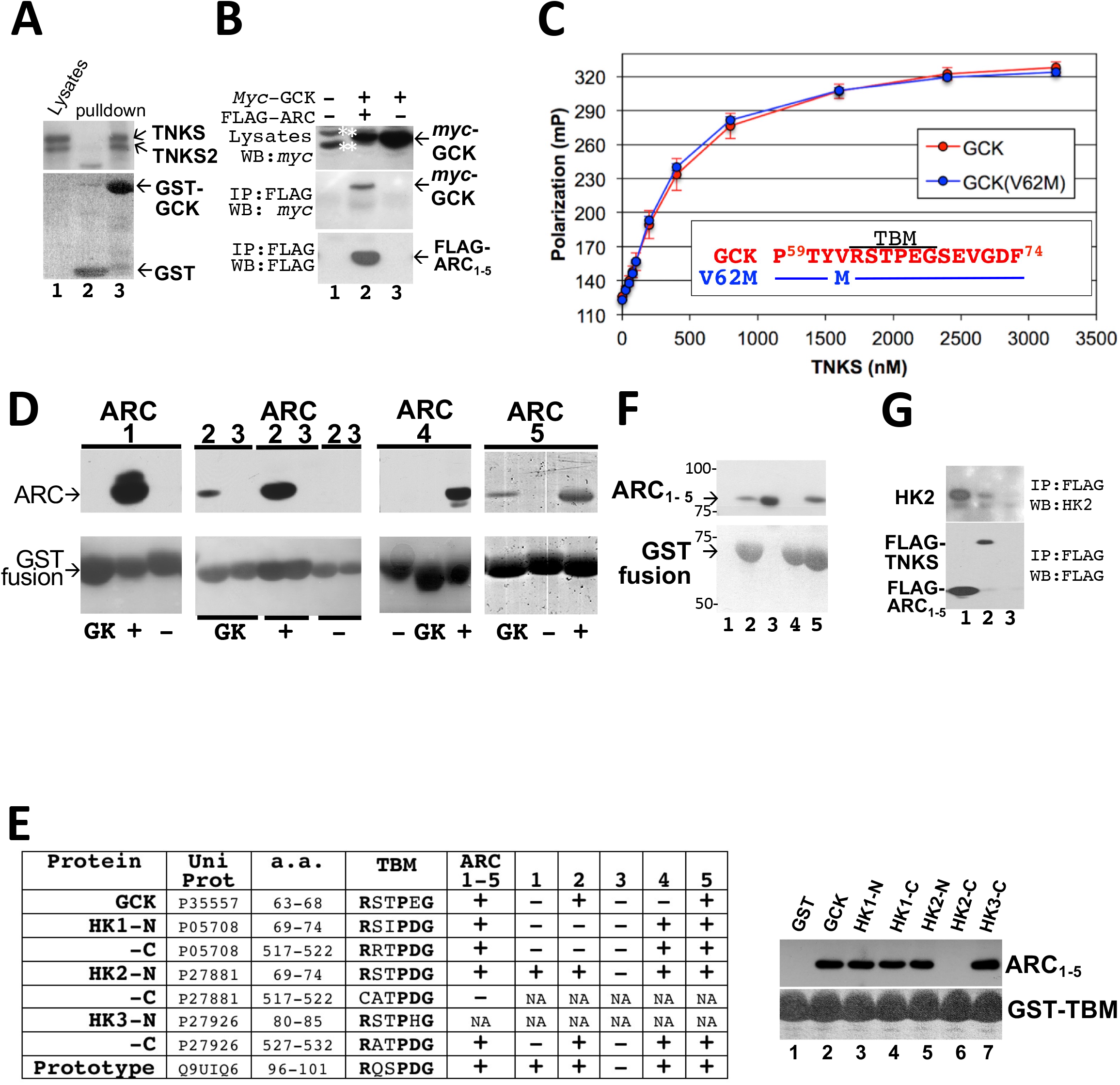
Interaction of TNKS with glucokinase, HK2, and the TBMs of HK1 and HK3. **A.** Fibroblast lysates (lane 1) were incubated with resins containing GST (lane 2) or GST-GCK fusion proteins (lane 3). The lysates and the GST pulldown were immunoblotted for TNKS (upper panel) and stained for GST fusions (lower panel). **B.** HEK293 cells were transfected with *myc-GCK* and FLAG-ARC_1-5_ vectors as indicated. Cell lysates were immunoprecipitated with anti-FLAG M2 agarose beads. The lysates (upper panel) and the immunoprecipitate (lower panels) were immunoblotted for *myc* (upper panels) and FLAG (lower panel). * indicates nonspecific bands flanking *myc*-GCK. **C.** Fluorescence polarization analysis of binding between TNKS ARC_1-5_ and a fluorophore-conjugated 16-mer peptide representing wild-type GCK (in red) or V62M-GCK (in blue). The data points and the error bars represent the averages and SD of 3 independent experiments. The lines represent the fit of a two-state binding model to the data. The TBM is marked by a line above the sequence. **D.** Individual ARCs of TNKS purified as a (His)_6_-SUMO fusions were incubated with resins containing GST alone (marked as -), GST fused to the prototypical TBM RQSPDG (marked as +), or the TBM of GCK (marked as GK). Resin-captured proteins were immunoblotted with an anti-His-tag antibody (upper panels). GST fusions were stained using Coomassie blue (lower panels). **E.** TBM hexapeptides from indicated hexokinases were examined for interaction with ARC_1-5_ and individual ARCs of TNKS as described in panel **D**. NA: not determined. The anti-His-tag immunoblot shows pulldown of (His)_6_-ARC_1-5_ (upper panel) by GST-TBM (lower panel). **F.** Purified ARC_1-5_ was incubated with resins containing GST (lane 1), GST fused to HK2-N, HK2-C or GCK (lanes 2, 4, and 5), or GST fused to the TBM prototype RQSPDG (lane 3). Resin-captured proteins were separated by PAGE and immunoblotted with an anti-His-tag antibody (upper panel) or stained with Ponceau S (lower panel). GST and GST-TBM (lanes 1 and 3) ran off the gel. **G.** HEK293 cells were transfected with an HK2 vector in combination with FLAG-ARC_1-5_ (lane 1), FLAG-TNKS (lane 2), or an empty vector (lane 3). Extracts were immunoprecipitated with anti-FLAG M2 affinity resins, and the precipitate was immunoblotted for HK2 (upper panel) or FLAG (lower panel).

### The N-terminal half of HK2 also contains a TBM and interacts with TNKS

The hinge region of the N-half of HK2 contains a hexapeptide (RSTPDG, aa 69-74) that matches the consensus TBM (RxxPDG). Indeed, both HK2-N and its TBM (**Fig. 2F**) pulled down ARC_1-5_, and ARC-1, 2, 4, and 5 was each sufficient to mediate the interaction **(Figs. 2E-F**). Full-length HK2 co-immunoprecipitated with TNKS and ARC_1-5_ (**Fig. 2G**). By contrast, the candidate TBM in HK2-C (CATPDG, aa 517-522) lacks the critical Arg at position 1 of the motif (**Fig. 2E**), and no interaction with ARC_1-5_ was detected **(Figs. 2E-F**). We did not examine HK1 or HK3 for potential interaction with TNKS. However, their TBMs did interact with ARC_1-5_ and 2 or 3 individual ARCs of TNKS (**Fig. 2E**). Our data therefore suggest that like GCK, all hexokinases use a conserved TBM in the hinge region to interact with TNKS.

### The GCK conformational selectivity of TNKS

Among the pathogenic mutations in the allosteric site of human GCK, V62M at position −1 of the TBM causes diabetes, whereas substitutions at positions 2, 3 and 6 (S64Y, T65I, G68V and G68D) of the TBM cause congenital hypoglycemia. We investigated the impact of these mutations on TNKS binding. In accordance with the TBM sequence rules established using a peptide library [33], G68V and G68D at the critical position 6 effectively abolished TNKS binding (**Fig. 3A**), whereas T65I (substituting position 3 with a less favorable residue) attenuated the binding by >4 fold (**Fig. 3B**). V62M at the ostensibly silent position −1 [33] was neutral to TNKS binding when tested in a GCK peptide (**Fig. 2C**) but robustly augmented the binding of full-length GCK (**Fig. 3C**). Conversely, S64Y at position 2, although predicted to be neutral by the TBM sequence rules [33], actually attenuated GCK binding to TNKS by ~4 fold (**Fig. 3B**). To account for how V62M and S64Y might violate the TBM sequence rules, we used the conformational selection model to hypothesize that only a select GCK conformer can present its TBM in a configuration that is favorable to engaging TNKS, and that this configuration was enriched by V62M and depleted by S64Y to robustly impact TNKS recruitment.

**Figure 3.**
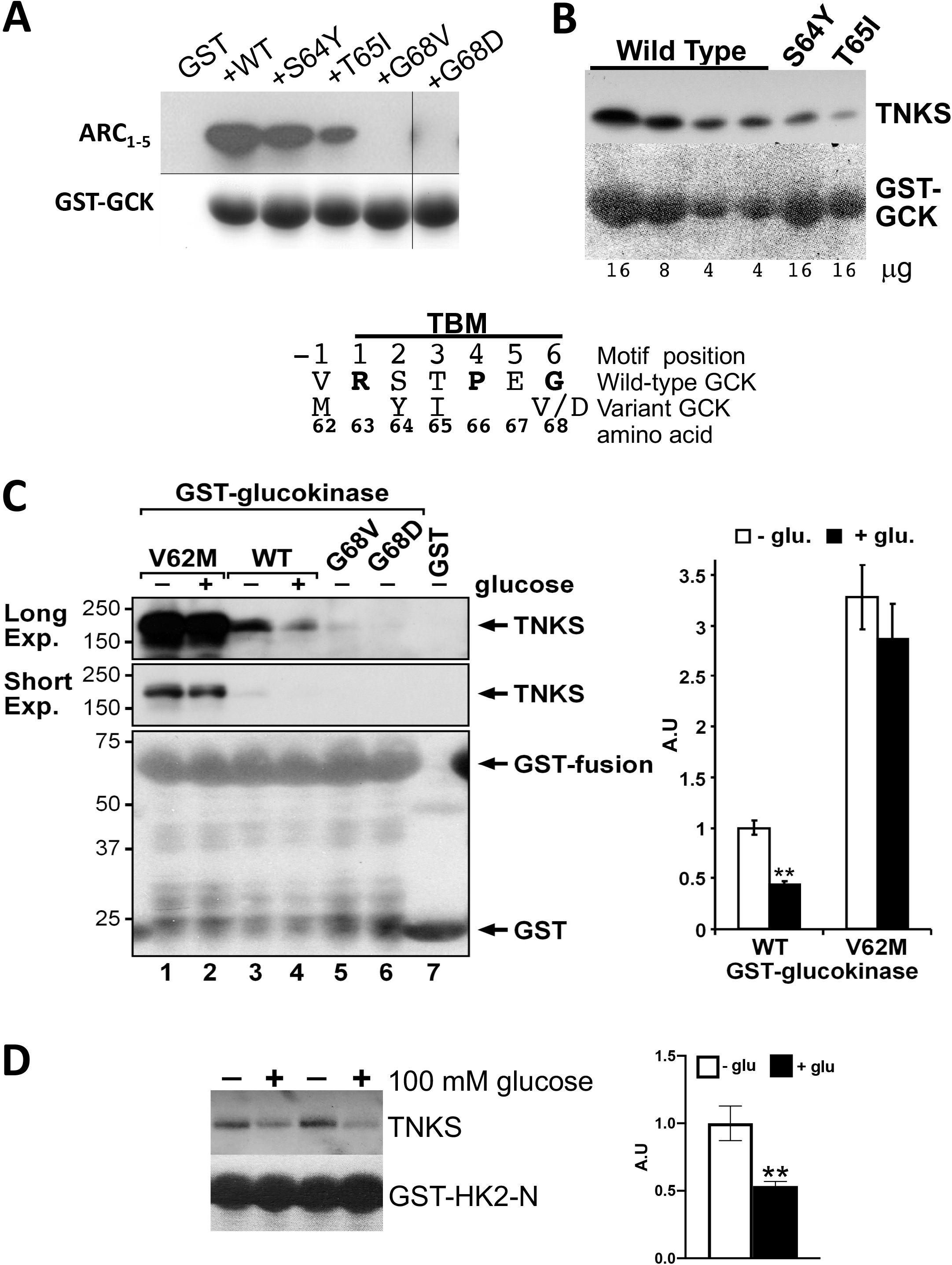
Effects of glucose and pathogenic mutations on GCK and HK2 interaction with TNKS. **A.** Purified ARC_1-5_ was incubated with resins containing GST control (the first lane) or GST fusions of wildtype GCK or the indicated variants (the remaining lanes). The pulldown was immunoblotted with anti-His-tag antibodies (upper panel) and stained for GST fusions (lower panel). **B.** Fibroblast lysates were incubated with resins containing GST-fusion of wild-type GCK (16, 8, or 4 μg protein as indicated), the S64Y variant (16 μg), or the T65I variant (16 μg). The GST pulldown was immunoblotted for TNKS (upper panel). GST fusions were visualized by Ponceau staining (lower panel). **C.** Fibroblast lysates were incubated with resins containing GST fusions of wild-type GCK or the indicated variants in the presence of 0 mM or 100 mM glucose as indicated. The pulldown was immunoblotted for TNKS (upper panel) and quantified by densitometry. Bar graph shows the efficiency of TNKS pulldown, normalized to GST-GCK and 0 mM glucose (corresponding to lane 3) as 1.0. **D**. Fibroblast lysates were incubated with resins containing GST fused to the N-terminal half of HK2 in the presence of 0 mM or 100 mM glucose as indicated. The GST pulldown was immunoblotted for TNKS (upper panel) and Ponceau-stained for GST-HK2-N (lower panel). The bar graph shows the efficiency of the pulldown as in **D**, normalized to 0 mM glucose as 1.0, and represents the average of 3 independent experiments, each conducted in >5 replicates.

### Glucose inhibits GCK interaction with TNKS

The GCK conformational selectivity further predicts that glucose, by shifting GCK toward the closed form [5, 6], might impact TNKS binding. Indeed, **Fig. 3C** shows that glucose (100 mM) robustly diminished TNKS pulldown by GST-GCK, indicating that the closed (most glucose-avid) form is not the putative TNKS-avid GCK conformer. Since the putative conformer appeared to be enriched in the V62M variant (**Fig. 3C**), we expected the V62M-TNKS complex to be less sensitive to the destabilizing effect of glucose. Indeed, **Fig. 3C** shows that TNKS pulldown by V62M was largely glucose insensitive.

The observed glucose effect on wild-type GCK was consistent with TNKS’s selection for any of the glucose-averse conformers: open, wide-open, or super-open GCK. Among these, only the open conformer is accessible to HK2 too. We therefore reasoned that if glucose similarly inhibits the HK2-TNKS interaction, then the open conformation can be designated as TNKS-avid. Indeed, **Fig. 3D** shows that glucose robustly attenuated TNKS pulldown by HK2 (N-half). Collectively, these findings suggested that TNKS selected for the open conformer shared by HK2 and GCK, and that glucose inhibited TNKS binding by depopulating this conformer to enrich the active conformer. As a precedent for this scenario, the glucokinase regulatory protein GKRP selectively interacts with the super-open (glucose-averse) GCK conformer, and glucose opposes this interaction [35, 36].

### TNKS inhibits the catalytic activity of GCK

GKRP, by trapping GCK in the super-open form, inhibits its kinase activity [36, 37]. By analogy, TNKS’s binding to the open GCK should similarly inhibit glucose phosphorylation by depopulating the active conformer. This was tested by determining the kinase activity of purified GST-GCK in a colorimetric assay, where glucose conversion to glucose-6-phosphoate (G6P) was coupled by the dehydrogenase G6PD to the production of chromogenic NADPH. **Fig. 4A** shows that pre-incubation with TNKS ARC_1-5_ (5x molar excess, at 0 mM glucose) inhibited GCK catalysis by ~40% at 100 mM glucose. Inhibition by ARC_1-5_ was also observed at lower glucose levels (**Fig. 4B**). Regardless of ARC pre-incubation, phosphorylation by GCK at 8 mM glucose was ~50% compared to at 100 mM (**Fig. 4C**).

**Figure 4.**
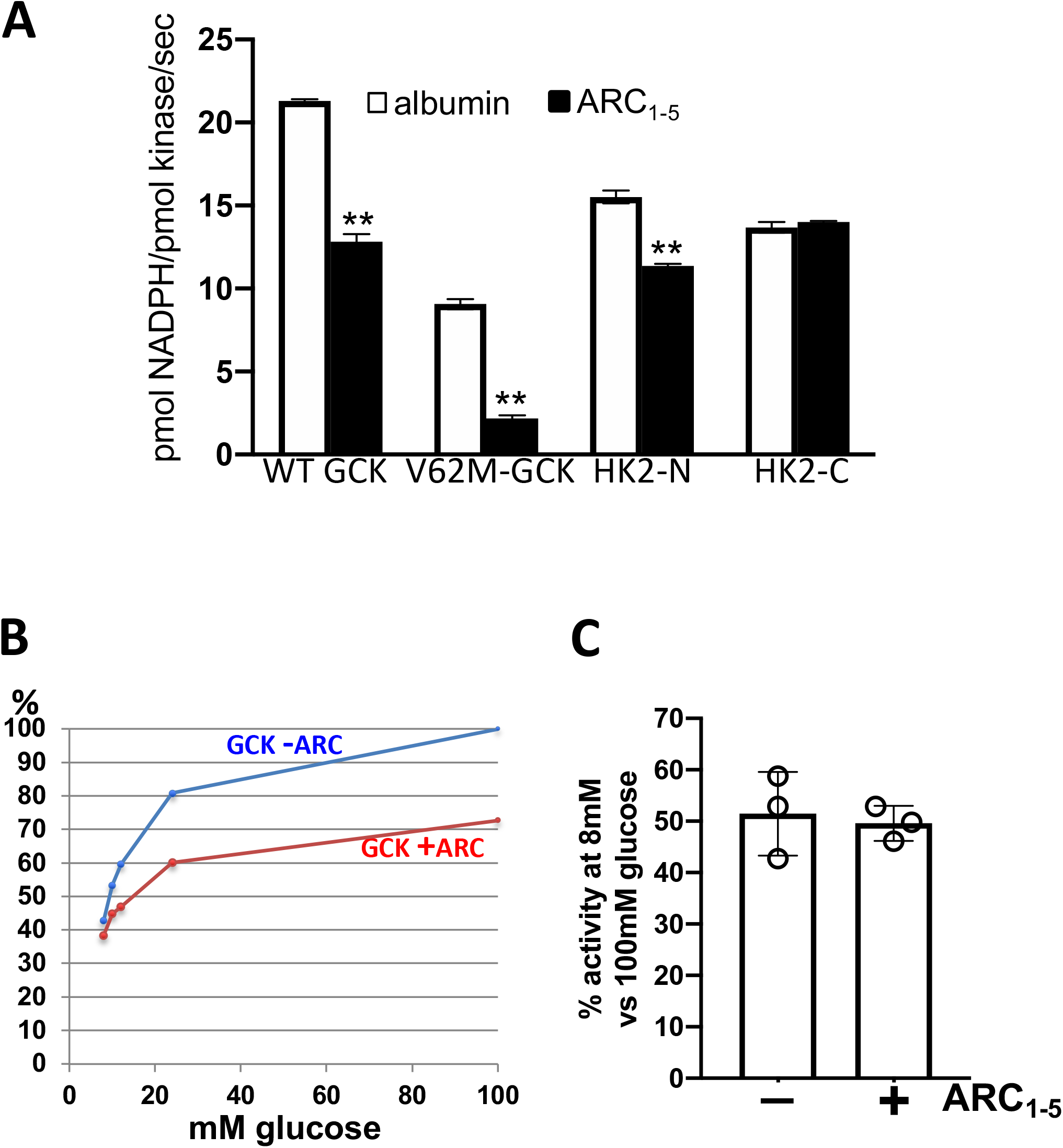
Effect of TNKS on the kinase activity of GCK (wild type and V62M), HK2-N, and HK2-C. **A.** GST fusions of the indicated kinase were pre-incubated with ARC_1-5_ or albumin. Kinase activity at 100 mM glucose was determined at 30 °C in triplicates as described in Materials and Methods. **B.** The kinase activity of wild-type GST-GCK, with or without pre-incubation with ARC_1-5_, was determined as in **A** but in the presence of 8, 10, 12, 24, or 100 mM glucose, and normalized to 100 mM glucose without ARC as 100%. **C.** The kinase activity of wild-type GST-GCK, with and without ARC pre-incubation, was determined at 8 mM and 100 mM glucose as in **A**. Activity ratios between the two glucose concentrations (8 mM over 100 mM) were averaged from 3 independent experiments.

To confirm that the observed inhibition in our kinetic assay was not mediated by impurities in our ARC preparations, we found that the inhibition was abrogated using pre-adsorption over GST-TBM resins, but not GST control, to deplete ARC_1-5_ from our ARC preparations (data not shown). To confirm that ARC_1-5_ inhibited GCK instead of G6PD in our coupled assays, we found that ARC did not affect assay readout if GCK was replaced by its product G6P (data not shown).

Compared to wild-type GCK, the greater affinity of the V62M variant for TNKS **(Fig. 3C)** suggested that it should be more sensitive to ARC inhibition. This was indeed the case (**Fig. 4A,** 76% inhibition of V62M *vs*. 40% for WT). As for hexokinase II, **Fig. 4A** shows that ARC inhibited HK2-N catalysis but not HK2-C, in line with selective binding to HK2-N (**Fig. 2F**). Collectively, our kinetic assays suggested that for HK2-N, GCK, and particularly the V62M variant, TNKS binding trapped the kinases in a catalytically inactive conformation.

Although our kinetic assays compared various kinases in parallel for sensitivity to TNKS inhibition (**Fig. 4A**), they were not intended for comparing the intrinsic activities of these kinases. This is because the kinases were not purified in parallel and might have undergone different extents of denaturation. In support of this caveat, V62M is more prone than wild-type GCK to thermal and oxidative inactivation [38, 39].

### Effects of TNKS knockdown on the two regulatory pathways of insulin secretion

Since TNKS inhibited GCK *in vitro*, we predicted knockdown of TNKS would phenocopy the pharmacological effects of GCK activators (GKAs) on the triggering pathway of insulin secretion, namely augmenting the response to 3-14 mM of glucose [14, 40]. Indeed, upon transfecting MIN6 cells with TNKS-specific dicersubstrate dsiRNA-A or -B (**Fig. 5A**) or a conventional siRNA (**Fig. 5B**), insulin secretory response to 2-8 mM glucose exceeded mock-transfected cells. Since the augmentation by GKAs is lost at >20 mM glucose [14, 40], we had expected TNKS knockdown to be neutral to insulin secretion at >20 mM glucose. However, the knockdown actually suppressed insulin secretion at 22 and 25 mM glucose despite expanding intracellular insulin stores (**Figs. 5B and C**). A similar inhibition was observed when TNKS-knockdown cells were stimulated by the GCK-independent secretagogue pyruvate, suggesting that the inhibition was not reflective of the triggering pathway.

**Figure 5.**
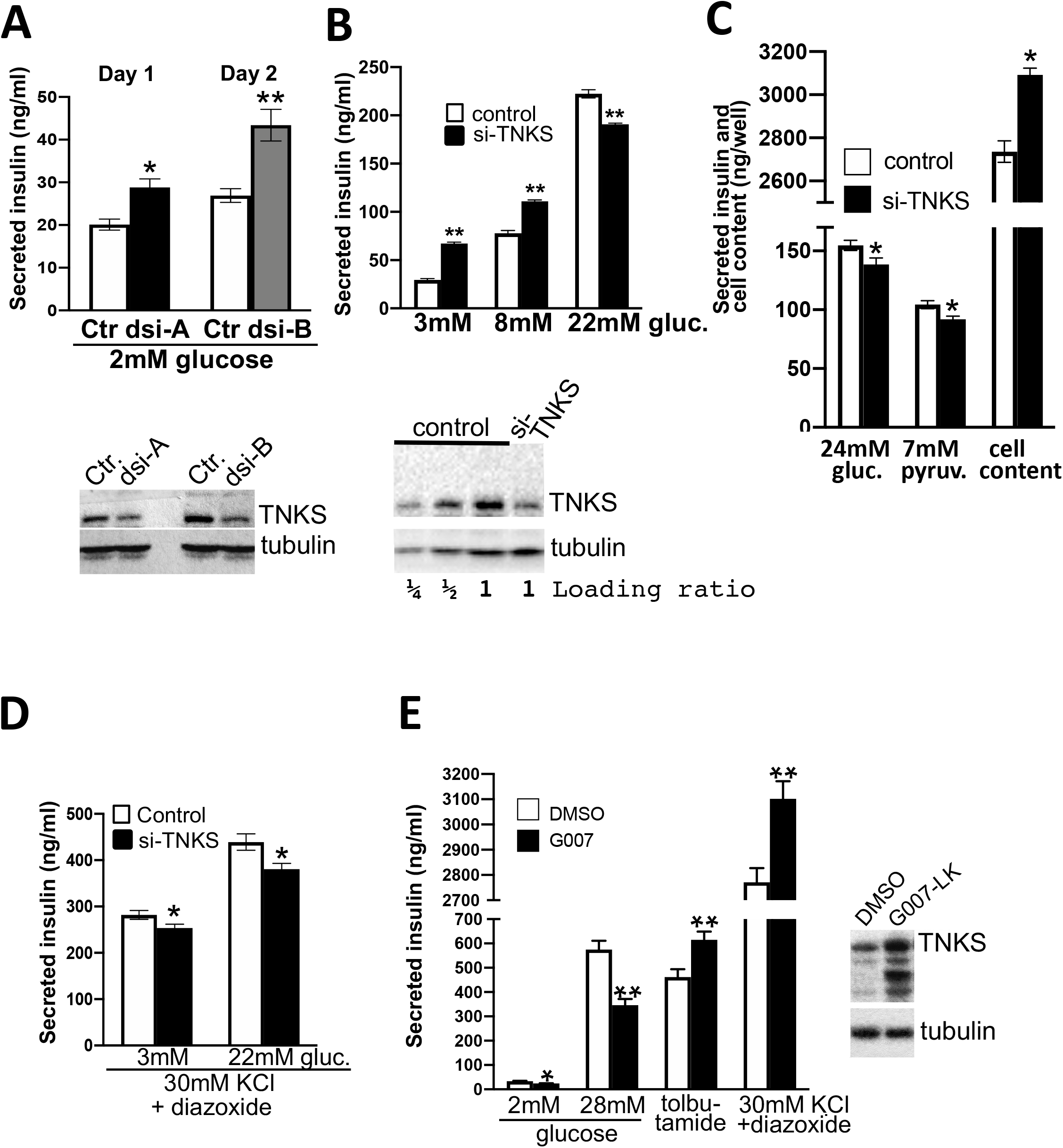
Effects of TNKS knockdown and TNKS inhibitor on insulin secretion in MIN6 cells. **A.** MIN6 cells were transfected with vehicle control (Ctr) or dicer-substrate TNKS dsiRNA-A, -B. After one (dsi-A) or 2 days (dsi-B), insulin secretion over 1 hr at 2 mM glucose was determined. Cell lysates were immunoblotted for TNKS and tubulin as described in Materials and Methods. **B.** MIN6 cells were transfected with a conventional TNKS siRNA or vehicle control. The next day, insulin secretion over 1 hr of stimulation with the indicated glucose concentration was determined. Cell lysates at the indicated loading ratio were immunoblotted for TNKS and tubulin to estimate the knockdown efficiency as between 50-75 %. **C.** MIN6 cells were transfected as in **B**. The insulin amount secreted during a 1-hr stimulation with 25 mM glucose or 7 mM pyruvate, and the cellular insulin content after glucose stimulation were determined. **D.** MIN6 cells were transfected as in **B**. After stimulation with 30 mM KCl and 100 μM diazoxide (a K_ATP_ channel opener that maximizes KCl effect) in combination with 3 mM or 27 mM glucose for 1 hr, insulin concentration in conditioned media was determined. **E.** Seven days past confluence, MIN6 cells were treated with DMSO or 2 μM G007-LK in culture media for 7 hr. Cells were then treated for 1 hr in KRB containing 2 mM glucose (with DMSO or G007-LK) before stimulation for another hr in KRB (with DMSO or G007-LK) containing glucose (2 mM or 28 mM), 300 μM tolbutamide, or 30 mM KCl plus 100 μM diazoxide. Conditioned media were analyzed for insulin concentration, and cell lysates were immunoblotted for TNKS and tubulin. **A-E:** Each bar represents the average (+SEM) of 6 wells except for 2 wells at 2 mM glucose in panel **E.**

To implicate the amplifying pathway instead, **Fig. 5D** shows that the inhibition persisted when 30 mM KCl was used to selectively unmask the amplifying pathway by bypassing the triggering pathway. Collectively, our findings indicated that TNKS knockdown augmented the triggering pathway as expected but also attenuated the amplifying pathway. Between these two opposing effects, the triggering pathway dominated the overall TNKS impact on the insulin response to physiological (<10 mM) glucose levels **(Figs. 5A-B)**.

### Effects of pharmacological TNKS stabilization are opposite to TNKS knockdown

To complement the knockdown approach, we treated MIN6 cells with the TNKS-specific PARsylation inhibitor G007-LK [41]. By blocking PARsylation-driven degradation, G007-LK increased TNKS abundance albeit in a catalytically inactive state. **Fig. 5E** shows that G007-LK impaired glucose-stimulated insulin secretion, consistent with inhibition of GCK glucose sensing by excess TNKS. By contrast, in cells treated with KCl or the K_ATP_ channel blocker tolbutamide to selectively unmask the amplifying pathway, which modulates the amplitude of insulin vesicle exocytosis in response to cytosolic calcium surges [12, 42], G007-LK augmented insulin secretion **(Fig. 5E**) without affecting cellular insulin stores (data not shown).

Collectively, our data indicate that G007-LK inhibited the triggering pathway as expected from TNKS excess but also augmented the amplifying pathway of insulin secretion. Neither effect was attributable directly to inhibition of TNKS-mediated PARsylation *per se*, since both G007-LK and TNKS knockdown diminished the PARsylation whereas their opposite impacts on insulin secretion tracked with opposite changes in TNKS abundance. We therefore reasoned that both experimental interventions acted by changing TNKS abundance, not its catalytic activity.

## DISCUSSION

Tankyrase is a molecular scaffold that has been implicated in diverse cellular processes. We were the first to identify RxxPDG as its binding motif and to delineate 5 ankyrin-repeat clusters (ARCs) in TNKS that recruit this motif [23, 43]. In TNKS-deficient mice, we found decreased blood glucose levels and excessive insulin secretion [27]. This pancreatic beta-cell phenotype prompted us to investigate whether TNKS impacts glucose sensing by GCK, using HK2 as a control.

We found that GCK and HK2 (N-terminal half) use their respective TBM to interact with multiple ARCs in TNKS (**Fig. 2**). For both kinases, glucose inhibits their TNKS interaction (**Fig. 3**) and, conversely, the interaction inhibits glucose phosphorylation (**Fig. 4**). When GCK was mutagenized to harbor the diabetogenic mutation V62M preceding the TBM (aa 63-68), the interaction with TNKS became more avid and glucose resistant (**Fig. 3C**). We attributed this effect to a redistribution of GCK conformations that enhances TBM recruitment by TNKS, since V62M was neutral to the recruitment when examined as a peptide (**Fig. 2C**). To account for our data, we propose that TNKS selectively binds to the open (inactive) form of both GCK and HK2-N (**Fig. 6**). This conformational selectivity can explain why glucose, known to favor the closed (active) form of both kinases, attenuates their interaction with TNKS (**Figs. 3C-D**), and conversely why the interaction inhibits catalysis (**Fig. 4**).

**Figure 6.**
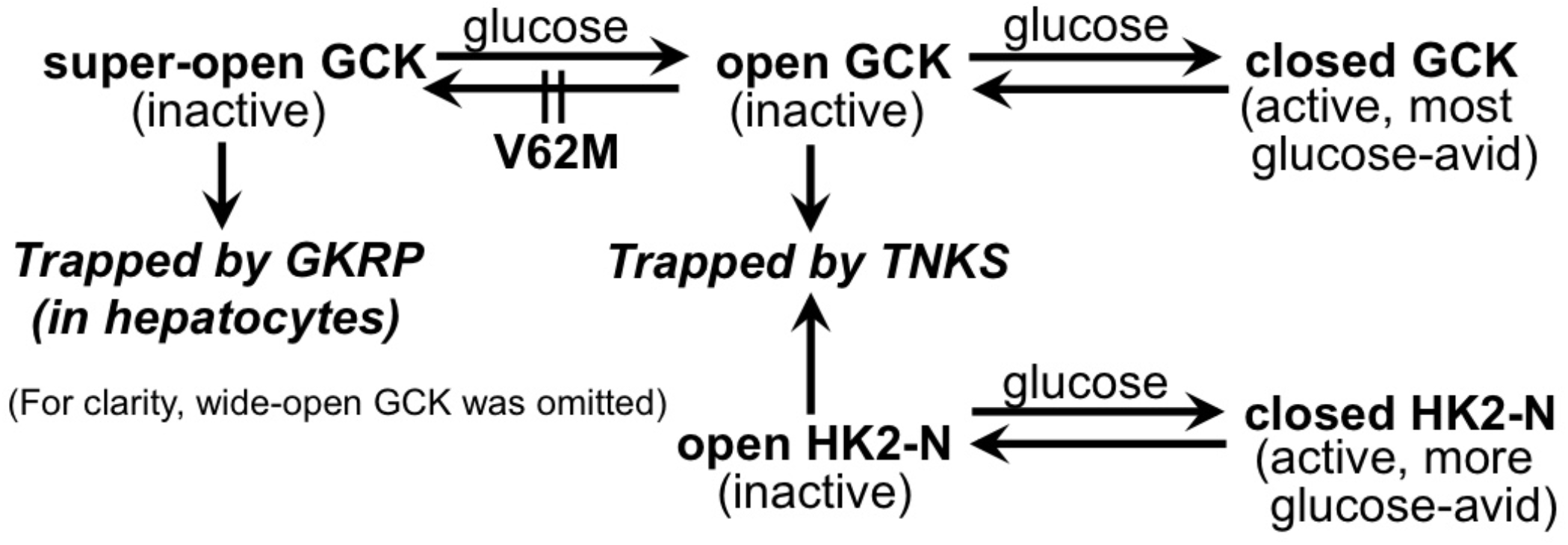
Proposed mechanism whereby TNKS selectively binds to the open (inactive) form of GCK and HK2, trapping them in an active state.

As a precedent for the proposed scenario, GKRP inhibits GCK catalysis by trapping it in the super-open form [36, 37], and this interaction is attenuated by glucose [35, 36]. Since V62M-GCK is resistant to GKRP recruitment [44] and inhibition [38], we further postulated that this mutation prevents GCK from populating the super-open form and thereby indirectly enriching the open (TNKS-avid) form. This would explain why V62M augments both GCK recruitment **(Fig. 3C)** and catalysis inhibition by TNKS (**Fig. 4A**).

The proposed model (**Fig. 6**) predicts that the structure of open GCK and particularly its TBM should be optimal for recruitment by TNKS. We therefore simulated docking of TNKS (ARC-2) with various GCK structures available in https://www.rcsb.org, including 1V4S, 3A0I and 3F9M (closed); 4DCH (open); and 1V4T (super-open). Surprisingly, every form exhibited significant steric clashes when docked with TNKS, and none of their TBMs followed the primarily linear backbone trajectory shared by other TNKS-bound TBM peptides (derived from Axin, TRF1, Mcl1, NuMA, 3BP2, and FNBP1 in https://www.rcsb.org). It therefore appears that the actual GCK conformation recruited by TNKS in solution is not represented in a straightforward manner by available crystal structures. We used open GCK as the best proxy in the model above to incorporate our protein binding and enzyme kinetic data.

It may seem paradoxical that 100 mM glucose inhibited GCK binding to TNKS (**Fig. 3C**) without protecting glucose phosphorylation from TNKS inhibition (**Figs. 4B-C**). A plausible explanation is that our kinetic assays pre-incubated GCK and TNKS (ARC_1-5_) in the absence of glucose to encourage protein complex formation. Upon adding 100 mM glucose to initiate the catalysis, the reaction rate was determined within 5-10 minutes, when glucose might not have had enough time to dissociate GCK from TNKS. Thus, a longer glucose exposure might have rescued GCK catalysis from inhibition by TNKS. Additional studies are required to determine how TNKS impacts GCK kinetic parameters (*V*_max_, *S*_0.5_, and glucose cooperativity).

Consistent with the inhibition of GCK by TNKS *in vitro*, our MIN6 data indicated that TNKS used its non-catalytic function to regulate insulin secretion, predominately by inhibiting the triggering pathway. In line with this view that TNKS does not use PARsylation to inhibit glucose sensing, G007-LK treatment did not increase the GCK content of MIN6 (data not shown) as would be expected of PARsylation substrates of TNKS [24, 45]. We also did not detect PARsylation of endogenous GCK in MIN6 or (*myc*)_2_-GCK co-transfected with TNKS in HEK293 cells (data not shown) using a published method [46].

In contrast to the triggering pathway of insulin secretion, our MIN6 data indicated that the non-catalytic function of TNKS augments the amplifying pathway, which modulates calcium-induced insulin vesicle exocytosis. In this regard, TNKS has been implicated in vesicular trafficking and exocytosis in adipocytes and myocytes [47–49]. In the beta cells, a potential tether for TNKS to regulate insulin vesicle exocytosis is VAMP8. This vesicle-associated membrane protein binds to TNKS [45] and is known to decorate insulin vesicles to promote their exocytosis [50, 51].

The demonstration of TNKS as an endogenous GCK inhibitor may account for the hyperinsulinemia and hypoglycemia of TNKS-deficient mice [27]. It may also help explain why V62M, a GCK mutation that enhances kinase activity *in vitro*, actually behaves like an inactivating mutation in humans [38]. This discordance could be due to V62M enhancing GCK binding and inhibition by TNKS **(Figs. 3C and 4A)**, an effect that would have eluded kinetic assays of isolated GCK.

The inhibition of GCK by TNKS may have metabolic implications beyond the beta cells. In pancreatic alpha cells, hepatocytes, and hypothalamic neurons, GCK also serves as a glucose sensor to regulate appetite and systemic glucose metabolism [52, 53]; these functions may be subject to TNKS regulation. More broadly, TNKS as a novel partner of HK2 **(Fig. 2G)** and an inhibitor of HK2-N **(Fig. 4A)** may modulate the glucose consumption of cancer cells. Specifically, overactive HK2 promotes tumor growth by increasing glucose consumption [4]. TNKS inhibitors that stabilize the TNKS protein in tumors may potentially curb tumor growth by inhibiting HK2.

## MATERIALS AND METHODS

### Plasmid vectors

A pcDNA3-based rat HK2 vector was a gift from Dr. John E. Wilson (Michigan State University; [54]). pGFP-GCK (human liver isoform [55]) was a gift from Dr. Rica Waterstradt (University of Rostock, Germany). A pcDNA3.1-based expression vector for (*myc*)_2_-GCK (human beta-cell isoform) was a gift from Dr. Ulupi Jhala (Univ. of California, San Diego). For bacterial expression as GST fusion proteins, the following were inserted into pGEX-4T1 between the *BamH1* and the *NotI* site: synthetic oligonucleotides encoding hexapeptide TBM followed by a stop codon; PCR-amplified cDNA encoding the N- or the C-terminal half of HK2 (aa 1-469; 470-917); and PCR-amplified cDNA encoding GCK. The latter was further subjected to site-directed mutagenesis using QuikChange Lightning kits (Agilent Inc.).

For bacterial expression with an N-terminal (His)_6_-SUMO tag (for affinity purification and greater solubility), individual ARCs of TNKS (aa 159-322, 318-475, 471-631, 627-790, and 797-983, respectively) were subcloned by PCR from FLAG-TNKS [28] and inserted into a pET28a-derived vector [56].

For CMV-driven expression, cDNA encoding TNKS ARC_1-5_ (aa 174-985) was subcloned by PCR into pFLAG-CMV_2_ (Addgene.org). All vectors subcloned using PCR were sequenced on both strands for verification.

### Protein purification

GST-TBM fusions were expressed and purified as described [28]. GST fusions of GCK, HK2-N, and HK2-C were similarly purified except that 50 mM glucose and 10 mM DTT were added to the lysis buffer prior to sonicating the *E. coli* pellet. Lysates from 1 liter of culture were centrifuged at 20,000 *g* for 20 min, and the supernatant was incubated with 300 μl glutathione-sepharose resins at 4 °C for 1-2 hours. The resins were washed 3 times with 50 mM Tris pH 8, 200 mM KCl, 50 mM glucose, and 10 mM DTT before elution using 15 mM glutathione. A typical yield was 250-500 μg proteins/per liter culture at 5 μg/μl.

TNKS ARC_1-5_ with a C-terminal (His)_6_ tag was overexpressed in the *E. coli* strain BL21(DE3)(Novogen Inc.) essentially as described [34] except that after elution from Ni-NTA agarose resins (Qiagen Inc.) by incubating with 400 mM imidazole for 1 hr at 4 °C, proteins were extensively dialyzed into 25 mM HEPES pH 8, 250 mM NaCl, and 1 mM DTT using Slide-A-Lyzer (Pierce Inc.) and concentrated to 10-12 mg/ml using Centricon 50 (Millipore Inc.). A typical yield was ~1 mg protein/liter culture. (His)_6_-SUMO fusions of individual ARCs were similarly expressed and purified from the *E. coli* strain Rosetta 2 (DE3)(Novagen Inc.).

### Protein interaction assays

GST pull-down assays were essentially as described [28]. Confluent 3T3-L1 fibroblasts were lysed by scraping the plates in buffer A (0.6 ml/15-cm plate)[28]. Soluble proteins were pre-cleared by incubating with glutathione resins containing GST (100 μg/plate). Supernatant (50 μl lysates per reaction) was incubated with resins containing 20 μg of GST-GCK or -HK2-N. The reactions were supplemented with 100 mM glucose when indicated. After overnight incubation at 4 °C, the resins were washed 3x in buffer A and captured proteins were analyzed by SDS-PAGE. GST fusions of TBM (10 μg proteins) were similarly used to pull down purified (His)_6_-SUMO fusions of individual ARCs (10 μg in 100 μl buffer A). Captured proteins were immunoblotted using an anti-His-tag antibody (clone 2A8, 1:1000, Abmart Inc.). GST fusions were stained with Ponceau S or 0.2% Coomassie Brilliant Blue (BIO-RAD Inc.). For co-transfection studies, calcium phosphate was used to transfect HEK293 cells with 2.5 μg of each plasmid per 6-cm plate. Cells were lysed 1-2 days later and the lysates were immunoprecipitated using anti-FLAG M2 affinity resins (10 μl per plate, Sigma Inc.). TNKS antibodies were as described (No. 465 in [57]; H-350 in [58]). HK2 antibody was from Cell Signaling Technology (#2106).

### Fluorescence polarization (FP) assay

Binding reactions were performed using 27 nM fluorophore-conjugated peptide and 0-3.2 μM TNKS ARC_1-5_ in 100 μL of a buffer containing 25 mM HEPES pH 8, 150 mM NaCl, 5.7 mM β-mercaptoethanol, 50 μg/mL BSA, and 5% glycerol. Mixtures were incubated at room temperature for 30 min and FP was measured on a Victor3V plate reader (PerkinElmer). A two-state binding model was fitted to the background-subtracted FP data using a quadratic formula in Sigma-Plot.

### Immunofluorescence

MIN6 cells grown on poly-lysine-coated coverslips were fixed in paraformaldehyde, permeabilized as described [28], and stained overnight for TNKS (mouse monoclonal E-10, Santa Cruz Inc., 5 μg/ml) and GCK (rabbit polyclonal, Aviva Inc. Cat. OAAF05763, 3 μg/ml). Similar staining patterns were observed (not shown) using two additional rabbit polyclonal GCK antibodies: one from Genetex Inc. (1:400) and the other a gift from Dr. Joan J. Guinovart (Univ. of Barcelona; 1:150). MIN6 cells grown in 24-well plates were also transfected with pGFP-GCK (1 μg DNA/well) using 2.5 μl Lipofectamine LTX and 1 μl PLUS reagent (Invitrogen Inc.). HEK293 cells grown in 24-well plates were transfected with pGFP-GCK (0.5 μg/well) using a calcium phosphate kit (Invitrogen Inc.). The next day, cells were fixed, permeabilized, and stained for TNKS as above and for β-tubulin (T2200, Sigma Inc. 0.6 μg/ml). Wide-field images were acquired using a Zeiss inverted microscope (Axio Observer Z1, Thornwood, NY).

### Enzyme kinetic assays

Stock preparations of GST-GCK/HK2 and (His)_6_-ARC_1-5_ were diluted into 20 μl of a binding buffer (50 mM HEPES pH 8, 25 mM KCl, 0.5% Triton, 0.1% β-mercaptoethanol, and 0.1% BSA) to the final concentrations of 0.5 μM (39 ng/μl) and 10 μM (880 ng/μl) respectively. As a negative control for (His)_6_-ARC_1-5_, albumin was used instead. After pre-incubation at 30 °C for 90 min to allow protein complex formation, coupled kinase-G6PD reactions were initiated by adding 980 μl of a pre-warmed cocktail that was adapted from [59, 60] to contain 100 mM glucose (unless otherwise specified), 5 mM ATP, 0.2 U/ml glucose-6-phosphate dehydrogenase (USB Corp.), 8 mM MgCl_2_, 0.4 mM NADP, 50 mM HEPES pH 8, 0.1% β-mercaptoethanol, 0.1% BSA, 0.5% Triton, 25 mM KCl, 0.5 mM EDTA, and 50 mM imidazole. Multiple aliquots of each reaction were loaded into 96-well plates (200 μl/well) and OD_340_ was recorded every 30 seconds for 15 min at 30 °C. Reaction rates were calculated after the OD_340_ slope became linear, typically between 5-10 min, and were corrected for background rate that was determined in parallel by leaving out the kinase from the reaction. OD_340_ was converted to NADPH concentrations using a molar extinction coefficient of 6,220 [61].

### Cell culture and treatments

MIN6 cells (a gift from Dr. Ulupi Jhala, Univ. of California, San Diego) were cultured in DMEM containing 100 mg/dL glucose, 4% FBS, 100 μM β-mercaptoethanol, and 0.15% NaHCO_3_, fed every 2-3 days, and seeded in 24-well plates (10^6^ cells per well) for experimentation. For G007-LK treatment, cells were allowed to grow past confluency for 5-7 days to optimize glucose responsiveness. Diazoxide and tolbutamide were from Sigma Inc. For TNKS knockdown, each well was transfected the day after seeding with 100 pmol dicer-substrate TNKS siRNA (27mer dsiRNA, Integrated DNA Technologies) or 21mer silencer select TNKS siRNA (ID s75314, ThermoFisher Scientific) using 7.6 μl X-tremeGENE (Roche Inc.) following the manufacturer’s instructions. Transfected cells were fed with the above culture media the next morning, and insulin secretion was assayed in the afternoon. The sequences of custom-made dsiRNAs are:

For insulin secretion assays, cells were rinsed and incubated in 300 μl/well of a KRB buffer [62] containing 1 or 2 mM glucose for 1 hr before stimulation with the indicated secretagogue in fresh KRB buffer (300 μl/well)

**Dsi-A:** 5’-GAGAUGCAGAGCACUAUUCGAGAGC-3’ 5’-GCUCUCGAAUAGUGCUCUGCAUCUCUU-3’
**Dsi-B:** 5’-GGAUGUUGUAGAACACUUGCUGCAG-3’ 5’-CUGCAGCAAGUGUUCUACAACAUCCUU-3’

for 1 hr. Conditioned media were assayed for insulin using an ELISA kit (Alpco Inc.). For cellular insulin content, cells were lysed in 300 μl/well of 0.25 N HCl and 80% ethanol, and insulin was extracted by shaking the plates vigorously on a shaker for 3 min. For immunoblotting, cells were lysed by vigorously shaking in SDS sample buffer (300 μl/well). Lysates were boiled before sheering by passing through small-gauge needles.

### Statistical analysis

Unpaired two-tailed *t*-test was performed for comparing between two groups. *p* value <0.05 was considered statistically significant. \**p* <0.05; \*\**p* <0.01. Data represent the mean ± SEM unless otherwise specified.

## ACKNOWLEDGMENTS

We are grateful to Dr. Wael M. Rabeh (New York University, Abu Dhabi) for insightful discussions. We thank Drs. John E. Wilson, Rica Waterstradt, Ulupi Jhala, and Joan J. Guinovart for generous gifts of reagents. We thank Drs. Joe Witztum, Len Deftos, Steven Chessler, and Chris Hupfeld for evaluable edits and comments.

